# Improved genome assembly and annotation for the rock pigeon (Columba livia)

**DOI:** 10.1101/220947

**Authors:** Carson Holt, Michael Campbell, David A. Keays, Nathaniel Edelman, Aurélie Kapusta, Emily Maclary, Eric Domyan, Alexander Suh, Wesley C. Warren, Mark Yandell, M. Thomas P. Gilbert, Michael D. Shapiro

**Affiliations:** Department of Human Genetics, University of Utah, Salt Lake City, UT, USA; USTAR Center for Genetic Discovery, University of Utah, Salt Lake City, UT, USA; Research Institute of Molecular Pathology, Vienna, Austria; Department of Biology, University of Utah, Salt Lake City, UT, USA; Department of Biology, Utah Valley University, Orem, UT, USA; Department of Evolutionary Biology (EBC), University of Uppsala, Uppsala, Sweden; Genome Institute at Washington University, St. Louis, MO, USA; Natural History Museum of Denmark, University of Copenhagen, Copenhagen, Denmark; Norwegian University of Science and Technology, University Museum, 7491 Trondheim, Norway

**Keywords:** *Columba livia*, rock pigeon, HiRise assembly, MAKER annotation

## Abstract

The domestic rock pigeon (*Columba livia*) is among the most widely distributed and phenotypically diverse avian species. This species is broadly studied in ecology, genetics, physiology, behavior, and evolutionary biology, and has recently emerged as a model for understanding the molecular basis of anatomical diversity, the magnetic sense, and other key aspects of avian biology. Here we report an update to the *C. livia* genome reference assembly and gene annotation dataset. Greatly increased scaffold lengths in the updated reference assembly, along with an updated annotation set, provide improved tools for evolutionary and functional genetic studies of the pigeon, and for comparative avian genomics in general.

## INTRODUCTION

Intensive selective breeding of the domestic rock pigeon (*Columba livia*) has resulted in more than 350 breeds that display extreme differences in morphology and behavior (Levi 1986; Domyan and Shapiro 2017). The large phenotypic differences among different breeds make them a useful model for studying the genetic basis of radical phenotypic changes, which are more typically found among different species rather than within a single species.

In genetic and genomic studies of *C. livia*, linkage analysis is important for identifying genotypes associated with specific phenotypic traits of interest (Domyan and Shapiro 2017); however, short scaffold sizes in the Cliv_1.0 draft reference assembly (Shapiro et al. 2013) hinder computationally-based comparative analyses. Short scaffolds also make it more difficult to identify structural changes, such as large insertions or deletions, that are responsible for traits of interest (Domyan et al. 2014; Kronenberg et al. 2015).

Here we present the Cliv_2.1 reference assembly and an updated gene annotation set. The new assembly greatly improves scaffold length over the previous draft reference assembly, and updated gene annotations show improved concordance with both transcriptome and protein homology evidence.

## MATERIALS & METHODS

### Genome sequencing and assembly

Genomic DNA from a female Danish tumbler pigeon (full sibling of the male bird used for the original Cliv_1.0 assembly (Shapiro et al. 2013)) was extracted from blood using a modified “salting out” protocol (Miller et al. 1988; modifications from http://www.protocol-online.org/prot/Protocols/Extraction-of-genomic-DNA-from-whole-blood-3171.html, accessed 06 February 2018)). Blood was frozen immediately after collection and stored at -80°C, and purified DNA was resuspended in 10 mM Tris-HCl. The sample went through 2 freeze-thaw cycles before being used to construct the libraries described below.

Extracted DNA was used to produce long-range sequencing libraries using the “Chicago” method (Putnam et al. 2016) by Dovetail Genomics (Santa Cruz, CA). Two Chicago libraries were prepared and sequenced on the Illumina HiSeq platform to a final physical coverage (1-50 kb pairs) of 390x.

Scaffolding was performed by Dovetail Genomics using HiRise assembly software and the Cliv_1.0 assembly as input. Briefly, Chicago reads were aligned to the input assembly to identify and mask repetitive regions, and then a likelihood model was applied to identify mis-joins and score prospective joins for scaffolding. The final assembly was then filtered for length and gaps according to NCBI submission specifications.

### Custom repeat library

A repeat library for *C. livia* was built by combining libraries from existing avian species (Zhang et al. 2014a) together with repeats identified *de novo* for the Cliv_2.1 assembly. *De novo* repeat identification was performed using RepeatScout (Price et al. 2005) with default parameters (>3 copies) to generate consensus repeat sequences. Identified repeats with greater than 90% sequence identity and a minimum overlap of 100 bp were assembled using Sequencher (Yokouchi et al. 1993). Repeats were classified into transposable element (TE) families using multiple lines of evidence, including homology to known elements, presence of terminal inverted repeats (TIRs), and detection of target site duplications (TSDs). Homology-based evidence was obtained using RepeatMasker (Smit et al. 1996), as well as the homology module of the TE classifying tool RepClass (Feschotte et al. 2009). RepClass was also used to identify signatures of transposable elements (TIRs, TSDs). We then eliminated non-TE repeats (simple repeats or gene families) using custom Perl scripts (available at https://github.com/4ureliek/ReannTE).

Our custom repeat analysis used the script ReannTE_FilterLow.pl consensus sequences as simple repeats or low complexity repeats if 80% of their length could be annotated as such by RepeatMasker (the library was masked with the option - noint). Next, we used the ReannTE_Filter-mRNA.pl script consensus sequences to RefSeq (Pruitt et al. 2007) mRNAs (as of March 7th 2016) with TBLASTX (Altschul et al. 1990). Sequences were eliminated from the library when: (i) the e-value of the hit was lower than 1E-10; (ii) the consensus sequence was not annotated as a TE; and (iii) the hit was not annotated as a transposase or an unclassified protein. The script ReannTE_MergeFasta.pl was then used to merge our library with a library combining RepeatModeler (Smit and Hubley 2008) outputs from 45 bird species (Kapusta et al. 2017) and complemented with additional avian TE annotations (International Chicken Genome Sequencing 2004; Warren et al. 2010; Bao et al. 2015). Merged outputs were manually inspected to remove redundancy, and all DNA and RTE class transposable elements were removed and replaced with manually curated consensus sequences, which were either newly (DNA elements) or previously generated (RTEs) (Suh et al. 2016).

### Repeat landscape

We used RepeatMasker software v4.0.7 (Smit et al. 2015) and our custom library to annotate the repeats in Cliv_2.1. RepeatMasker was run with the NCBI/RMBLAST v2.6.0+ search engine (-e ncbi), the sensitive (-s) option, the -a option in order to obtain the alignment file, and without RepeatMasker default libraries. We then used the parseRM.pl script v5.7 (available at https://github.com/4ureliek/Parsing-RepeatMasker-Outputs (Kapusta et al. 2017)), on the alignment files from Repeat Masker, with the -loption and a substitution rate of 0.002068 substitutions per site per million years (Zhang et al. 2014b). The script collects the percentage of divergence to the consensus for each TE fragment, after correction for higher mutation rate at CpG sites and the Kimura 2- Parameter divergence metric (provided in the alignment files from RepeatMasker). The percentage of divergence to the consensus is a proxy for age (the older the TE invasion, the more mutations will accumulate in TE fragments), to which the script applies the substitution rate in order to split TE fragments into bins of 1 My.

### Transcriptomics

RNA was extracted from adult tissues (brain, retina, subepidermis, cochlear duct, spleen, olfactory epithelium) of the racing homer breed, and one whole embryo each of a racing homer and a parlor roller (approximately embryonic stage 25 (Hamburger and Hamilton 1951)). RNA-seq libararies were prepared and sequenced using 100-bp paired-end sequencing on the Illumina HiSeq 2000 platform at the Research Institute of Molecular Pathology, Vienna (adult tissues), and the Genome Institute at Washington University, St. Louis (embryos). RNA-seq data generated for the Cliv_1.0 annotation were also downloaded from the NCBI public repository for *de novo* re-assembly. Accession numbers for these public data are SRR521357 (Danish tumbler heart), SRR521358 (Danish tumbler liver), SRR521359 (Oriental frill heart), SRR521360 (Oriental frill liver), SRR521361 (Racing homer heart), and SRR521362 (Racing homer liver).

Each FASTQ file was processed with FastQC (http://www.bioinformatics.babraham.ac.uk/projects/fastqc/) to assess quality. When FastQC reported overrepresentation of Illumina adapter sequences, we trimmed these sequences with fastx_clipper from the FASTX-Toolkit (http://hannonlab.cshl.edu/fastx_toolkit/). We used FASTX-Toolkit for two additional functions: runs of low quality bases at the start of reads were trimmed with fastx_trimmer when necessary (quality cutoff of -Q 33), and reads were then trimmed with fastq_quality_trimmer (-Q 33). Finally, each pair of sequence files was assembled with Trinity (Grabherr et al. 2011) version r20131110 using the --jaccard_clip option.

### Genome annotation

The pre-existing reference Gnomon (Souvorov et al. 2010) derived gene models for the Cliv_1.0 assembly (GCA_000337935.1) were mapped onto the updated Cliv_2.1 reference assembly using direct alignment of transcript FASTA entries. This was done using the alignment workflow of the genome annotation pipeline MAKER (Cantarel et al. 2008; Holt and Yandell 2011), which first seeds alignments using BLASTN (Altschul et al. 1990) and then polishes the alignments around splice sites using Exonerate (Slater and Birney 2005). Results were then filtered to remove alignments that had an overall match of less than 90% of the original model (match is calculated as percent identity multiplied by percent end-to-end coverage).

For final annotation, MAKER was allowed to identify *de novo* gene models that did not overlap the aligned Gnomon models. Protein evidence sets used by MAKER included annotated proteins from *Pterocles gutturalis* (yellow-throated sandgrouse) (Zhang et al. 2014a) and *Gallus gallus* (chicken) (International Chicken Genome Sequencing 2004) together with all proteins from the UniProt/Swiss-Prot database (Bairoch and Apweiler 2000; UniProt 2007). The transcriptome evidence sets for MAKER included Trinity mRNA-seq assemblies from multiple *C. livia* breeds and tissues (methods for transcriptome assembly are described above). Gene predictions were produced within MAKER by Augustus (Stanke and Waack 2003; Stanke et al. 2008). Augustus was trained using 1000 Cliv_1.0 Gnomon gene models that were split using the randomSplit.pl script into sets for training and evaluation. We followed a semi-automatic training protocol (https://vcru.wisc.edu/simonlab/bioinformatics/programs/augustus/docs/tutorial2015/training.html, accessed 9 February 2018). Repetitive elements in the genome were identified using the custom repeat library described above.

### Linkage map construction and anchoring to current assembly

Genotyping by sequencing (GBS) data was generated, trimmed, and filtered as previously described (Domyan et al. 2016). Reads were mapped to the Cliv_2.1 assembly using Bowtie2 (Langmead and Salzberg 2012). Genotypes were called using Stacks (Catchen et al. 2011), with a minimum read-depth cutoff of 10. Thresholds for automatic corrections were set using the parameters –min_hom_seqs 10, –min_het_seqs 0.01, –max_het_seqs 0.15. Sequencing coverage and genotyping rate varied between individuals, and birds with genotyping rates in the bottom 25% were excluded from map assembly.

Genetic map construction was performed using R/qtl v1.41-6 (www.rqtl.org) (Broman et al. 2003). For autosomal markers, markers showing segregation distortion (Chi-square, p < 0.01) were eliminated. Sex-linked scaffolds were assembled and ordered separately, due to differences in segregation pattern for the Z-chromosome. Z-linked scaffolds were identified by assessing sequence similarity and gene content between pigeon scaffolds and the Z-chromosome of the annotated chicken genome (Ensembl Gallus_gallus-5.0).

Pairwise recombination fractions were calculated for all autosomal and Z-linked markers. Missing data were imputed using “fill.geno” with the method “no_dbl_XO”. Duplicate markers were identified and removed. Within individual scaffolds, R/qtl functions “droponemarker” and “calc.errorlod” were used to assess genotyping error. Markers were removed if dropping the marker led to an increased LOD score, or if removing a non-terminal marker led to a decrease in length of >10 cM that was not supported by physical distance. Individual genotypes were removed if they showed with error LOD scores >5 (Lincoln and Lander 1992). Linkage groups were assembled from 2960 autosomal markers and 232 Z-linked markers using the parameters (max.rf 0.1, min.lod 6). In the rare instance that single scaffolds were split into multiple linkage groups, linkage groups were merged if supported by recombination fraction data; these instances typically reflected large physical gaps between markers on a single scaffold. Scaffolds in the same linkage group were manually ordered based on calculated recombination fractions and LOD scores.

To compare the linkage map to the original genome assembly (Cliv_1.0), each 90-bp locus containing a genetic marker was parsed from the Stacks output file “catalogXXX_tags.tsv” and queried to the Cliv_1.0 assembly using BLASTN (v2.6.0+) with the parameters –max_target_seqs 1 –max hsps 1. 3175 of the 3192 loci (99.47%) from the new assembly had a BLAST hit with an E-value < 4e-24 and were retained.

### Assembly comparisons

FASTA files from the Cliv_2.1 and colLiv2 (Damas et al. 2017) genome assemblies were hard masked using NCBI WindowMasker (Morgulis et al. 2006) and genome-wide alignments were calculated with LAST (Kielbasa et al. 2011). From these alignments, a genome-scale dotplot indicating syntenic regions was generated using SynMap (Lyons and Freeling 2008; Lyons et al. 2008).

The colLiv2 assembly is currently unannotated. Therefore, to compare gene content between assemblies, we estimated the number of annotated Cliv_2.1 genes absent from colLiv2 based on gene coordinates. Based on the length of LAST alignments, we calculated the percent of each Cliv_2.1 scaffold aligning to colLiv2. Scaffolds were divided into four groups based on alignments: Cliv_2.1 scaffolds that did not align to colLiv2, Cliv_2.1 scaffolds where LAST alignments to colLiv2 covered less than 50% of the total scaffold length, Cliv_2.1 scaffolds where LAST alignments to colLiv2 covered between 50% and 75% of the total scaffold length, and Cliv_2.1 scaffolds where LAST alignments to colLiv2 covered 75% or more of the total scaffold length. For each of these groups, the number of scaffolds containing genes was quantified. Many of these scaffolds are small, and some may be partially or completely missing from the alignment due to masking of repetitive elements. If annotated gene coordinates from Cliv_2.1 scaffolds fell partially or entirely within a region aligned to colLiv2, these genes were considered “present” in colLiv2. Thus, the number of genes marked as “absent” in colLiv2 might be a conservative estimate.

To compare the linkage map to colLiv2, each 90-bp locus containing a genetic marker was parsed from the Stacks output file “catalogXXX_tags.tsv” and queried to the colLiv2 assembly using BLASTN (v2.6.0+) with the parameters –max_target_seqs 1 –max hsps 1.

### Data availability

This Whole Genome Shotgun project has been deposited at DDBJ/ENA/GenBank under the accession AKCR00000000. The version described in this paper is version AKCR02000000. The Cliv_2.1 assembly, annotation, and associated data are available at ftp://ftp.ncbi.nlm.nih.gov/genomes/all/GCA/000/337/935/GCA_000337935.2_Cliv_2.1. RNA-seq data are deposited in the SRA database with the BioSample accession numbers SAMN07417936-SAMN07417943, and sequence accessions SRR5878849- SRR5878856. Assembly and RNA-seq data are publicly available in NCBI databases under BioProject PRJNA167554. File S1 contains Tables S1-S7. Files S2 and S3 contain recombination fraction data used to construct Figures 5a and 5b, respectively.

## RESULTS AND DISCUSSION

### Genome assembly

The final Cliv_2.1 reference assembly is 1,108,534,737 base pairs in length and consists of 15,057 scaffolds (Table 1). A total of 1,015 scaffolds contain a gene annotation. Completion analysis of the assembly using BUSCO v2 and the odb9 Vertebrata ortholog dataset (Simao et al. 2015) suggests that Cliv_2.1 is 72.9 (assembly) to 86.2% (annotation) complete. These statistics are nearly identical to the Cliv_1.0 assembly estimate of 72.3-86.4% (Table 2); therefore, we found no significant changes in completeness between the two assemblies. Because the Chicago libraries and HiRise assembly were designed to improve scaffolding of the original assembly, not to fill gaps, we did not expect substantial improvement to assembly completeness in Cliv_2.1. Instead, the major improvement to the Cliv_2.1 assembly is a substantial increase in scaffold length (Fig. 1a). Overall, The N50 scaffold length increased to 14.3 megabases, compared to 3.15 megabases for Cliv_1.0, a greater than 4-fold increase.

**Table 1.** Assembly statistics for Cliv_2.1.

|  |  |
| --- | --- |
| Estimated Physical Coverage | 389.7x |
| Total Length | 1,108,534,737 bp |
| Total scaffolds | 15,057 |
| Total scaffolds >1kb | 4,062 |
| Total scaffolds >10kb | 848 |

**Table 2.**
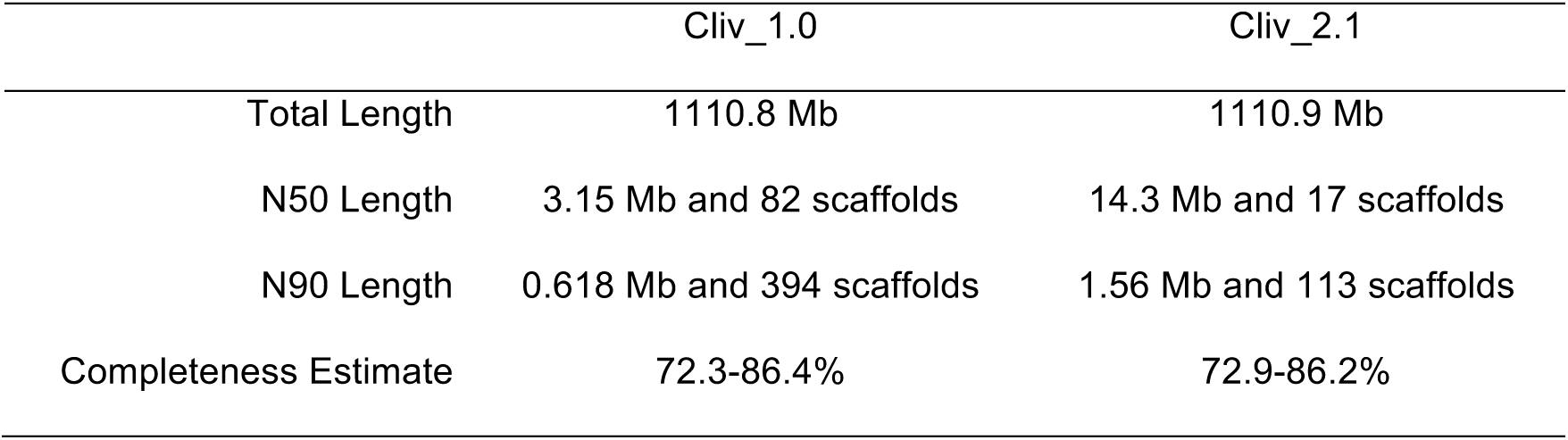
Assembly version comparison.

|  | Cliv_1.0 | Cliv_2.1 |
| --- | --- | --- |
| Total Length | 1110.8 Mb | 1110.9 Mb |
| N50 Length | 3.15 Mb and 82 scaffolds | 14.3 Mb and 17 scaffolds |
| N90 Length | 0.618 Mb and 394 scaffolds | 1.56 Mb and 113 scaffolds |
| Completeness Estimate | 72.3-86.4% | 72.9-86.2% |

**Figure 1.**
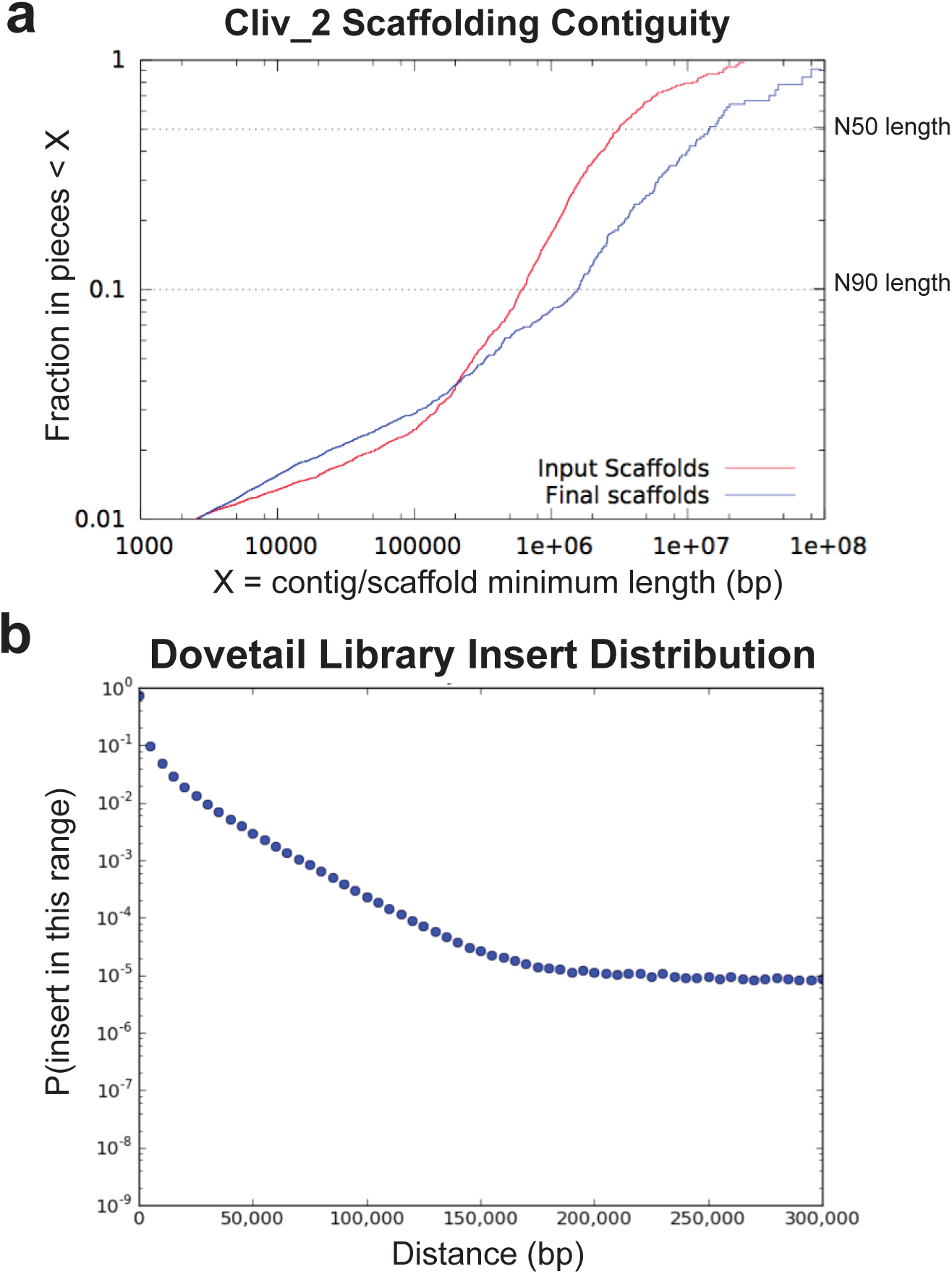
Assembly scaffolding contiguity and scaffolding library insert size distributions. (a) Scaffolding comparison between Cliv_1.0 (input scaffolds) and Cliv_2.1 (final scaffolds) assemblies. (b) Distribution of Dovetail Genomics “Chicago” library inserts.

The new assembly joins that, based on linkage mapping evidence (Domyan et al. 2016), we knew were physically adjacent but were still separated in Cliv_1.0 (see Table S1 for full catalog of positions of the original assembly in the new assembly, and Table S2 for full catalog of breaks in the original assembly to form the new assembly). For example, we previously determined that Cliv_1.0 Scaffolds 70 and 95 were joined based on genetic linkage data from a laboratory cross (Domyan et al. 2016). These two sequences are now joined into a single scaffold in the Cliv_2.1 assembly (see Table S6 for positions of genetic markers in Cliv_1.0 and Cliv_2.1). At least one gene model (RefSeq LOC102093126), which was previously split across two contigs, has now been unified into a single model on a single scaffold.

### Repeat landscape

Using our custom library, we identified 8.04% (89.1 Mb; Table S3) of the genome assembly as repeats, which is slightly higher than the previously published estimates of 7.25% (Zhang et al. 2014b) and 7.83% (Kapusta and Suh 2017). To illustrate the temporal dynamics of TE accumulation (see Methods), we split the amount of DNA of each TE class by bins of 1 million years (My) (Fig. 2). This landscape shows that TE accumulation has been consistent throughout time, with some potentially recently active elements. This includes CR1 LINEs (part of the non-LTR fraction), which are presumed to be inactive in most birds (Kapusta and Suh 2017), but comprise over 0.1 Mb of CR1 copies in the youngest bin (0-1 My) in the Cliv_2.1 assembly (Table S4).

**Figure 2.**
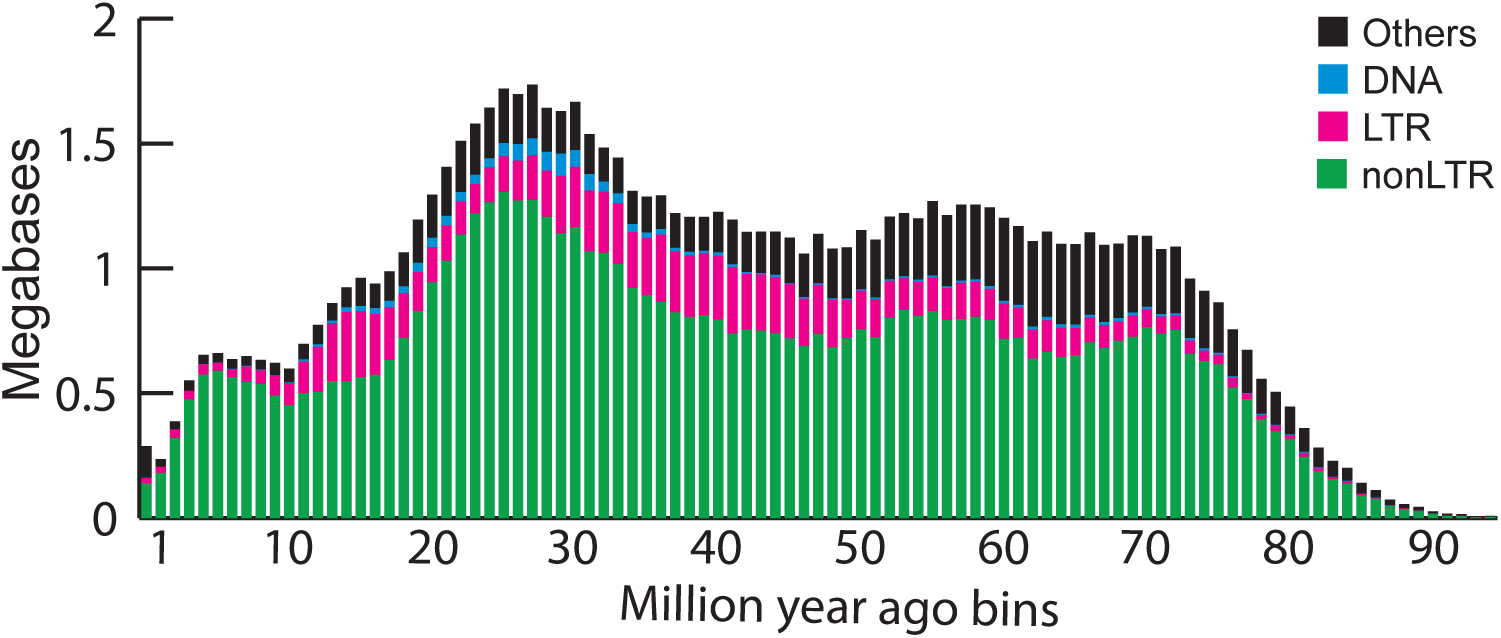
Temporal landscape of transposable elements. The amounts of DNA of each TE class were split into bins of 1 My, shown on the x axis (see Methods). We note that the lower detection of older elements (right of the graph) comes from a combination of lack of detection and TE removal, and that the amount of DNA corresponding to recent elements may be underestimated (recent copies are often collapsed in assemblies). The “Others” category primarily includes unclassified repeats.

### Transcriptome assemblies

A total of 1,936,543 transcripts were assembled from the 14 RNA-seq data sets. Numbers of assembled transcripts from each tissue are listed in Table 3. BUSCO analysis indicated 85.6% completeness of the union of transcriptome assemblies compared to the Vertebrata ortholog set.

**Table 3.**
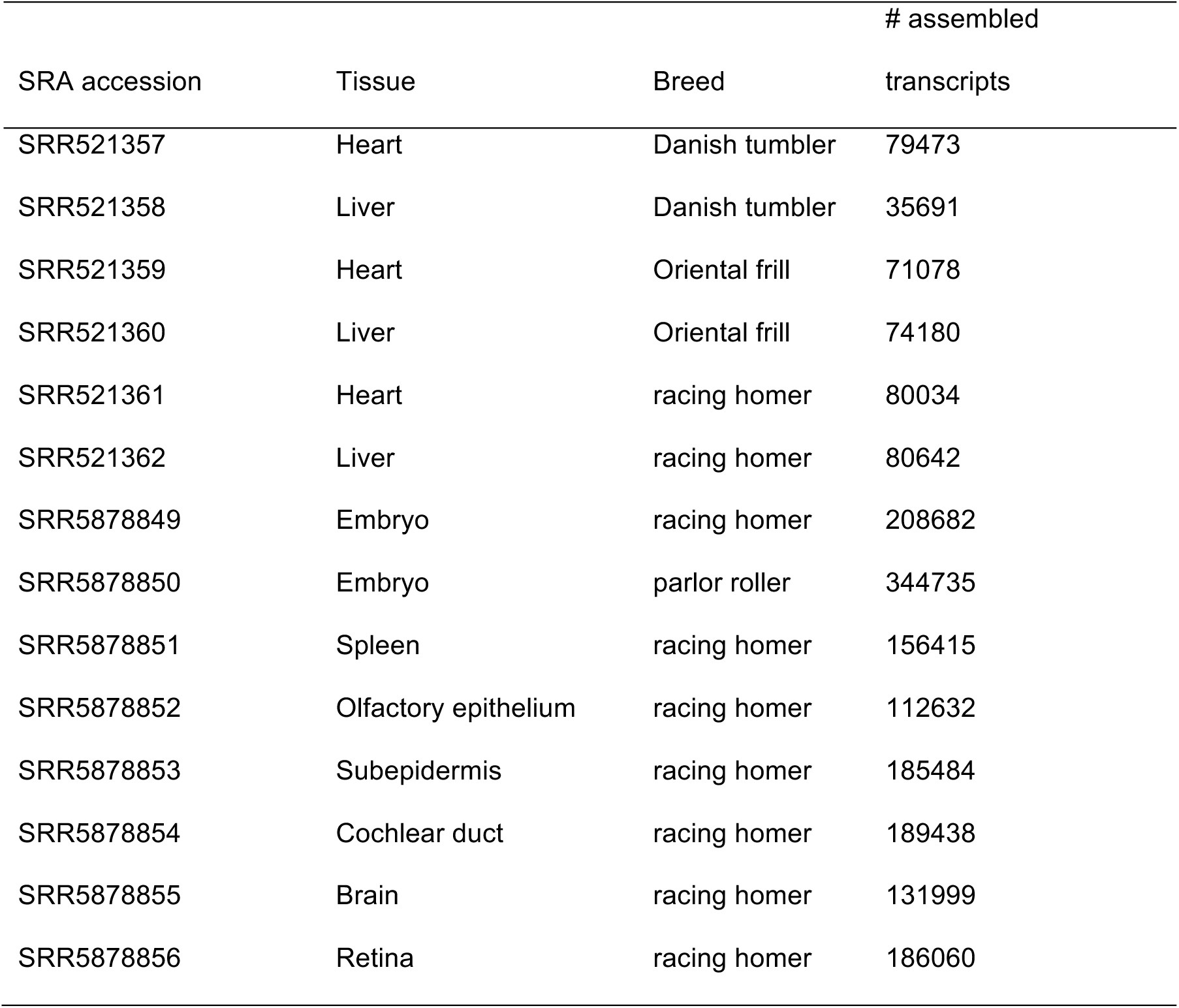
Transcriptome assembly summary.

| SRA accession | Tissue | Breed | # assembled |
| --- | --- | --- | --- |
|  |  |  | transcripts |
| SRR521357 | Heart | Danish tumbler | 79473 |
| SRR521358 | Liver | Danish tumbler | 35691 |
| SRR521359 | Heart | Oriental frill | 71078 |
| SRR521360 | Liver | Oriental frill | 74180 |
| SRR521361 | Heart | racing homer | 80034 |
| SRR521362 | Liver | racing homer | 80642 |
| SRR5878849 | Embryo | racing homer | 208682 |
| SRR5878850 | Embryo | parlor roller | 344735 |
| SRR5878851 | Spleen | racing homer | 156415 |
| SRR5878852 | Olfactory epithelium | racing homer | 112632 |
| SRR5878853 | Subepidermis | racing homer | 185484 |
| SRR5878854 | Cochlear duct | racing homer | 189438 |
| SRR5878855 | Brain | racing homer | 131999 |
| SRR5878856 | Retina | racing homer | 186060 |

### Annotation

The updated annotation set contains 15,392 gene models encoding 18,966 transcripts (Table 4). This represents a minor update of the reference annotation set as 94.7% of previous models were mapped forward nearly unmodified (90% exact match for 14,898 out of 15,724 previous gene models) and 494 new gene models were added to the Cliv_2.1 annotation set (Table 5).

**Table 4.** Annotation statistics for Cliv_2.1.

|  | Genes | Transcripts |
| --- | --- | --- |
| Total | 15,392 | 18,966 |
| match<sup>a</sup> | 14,898 | 18,472 |
| new | 494 | 494 |

<sup>a</sup> Count that match Cliv\_1.0 annotations with a value of at least 90% (match is calculated as % identity multiplied by % end-to-end coverage)

**Table 5.** Annotation version comparison.

|  | Cliv_1.0 | Cliv_2.1 |
| --- | --- | --- |
| Total Gene Models | 15,724 | 15,392 |
| *coding* | 15,022 | 14,683 |
| *non-coding* | 702 | 709 |
| Total Transcripts | 19,585 | 18,966 |
| *coding* | 18,569 | 18,148 |
| *non-coding* | 1016 | 818 |

The updated annotation set shows a modest improvement in concordance with aligned evidence datasets from mRNA-seq and cross species protein homology evidence relative to the Cliv_1.0 set as measured by Annotation Edit Distance (AED) (Eilbeck et al. 2009; Holt and Yandell 2011). As a result, transcript models in the Cliv_2.1 annotation tend to have lower AED values than the Cliv_1.0 set (Fig 3; the cumulative distribution function (CDF) curve is shifted to the left). Lower AED values indicate greater model concordance with aligned transcriptome and protein homology data. Furthermore, the Cliv_2.1 dataset displays greater transcript counts in every AED bin despite having slightly fewer transcripts overall compared to the Cliv_1.0 dataset (Table S5). The higher bin counts indicate that lower AED values are not solely a result of removing unsupported models from the annotation set, but rather suggest that evidence concordance has improved overall.

**Figure 3.**
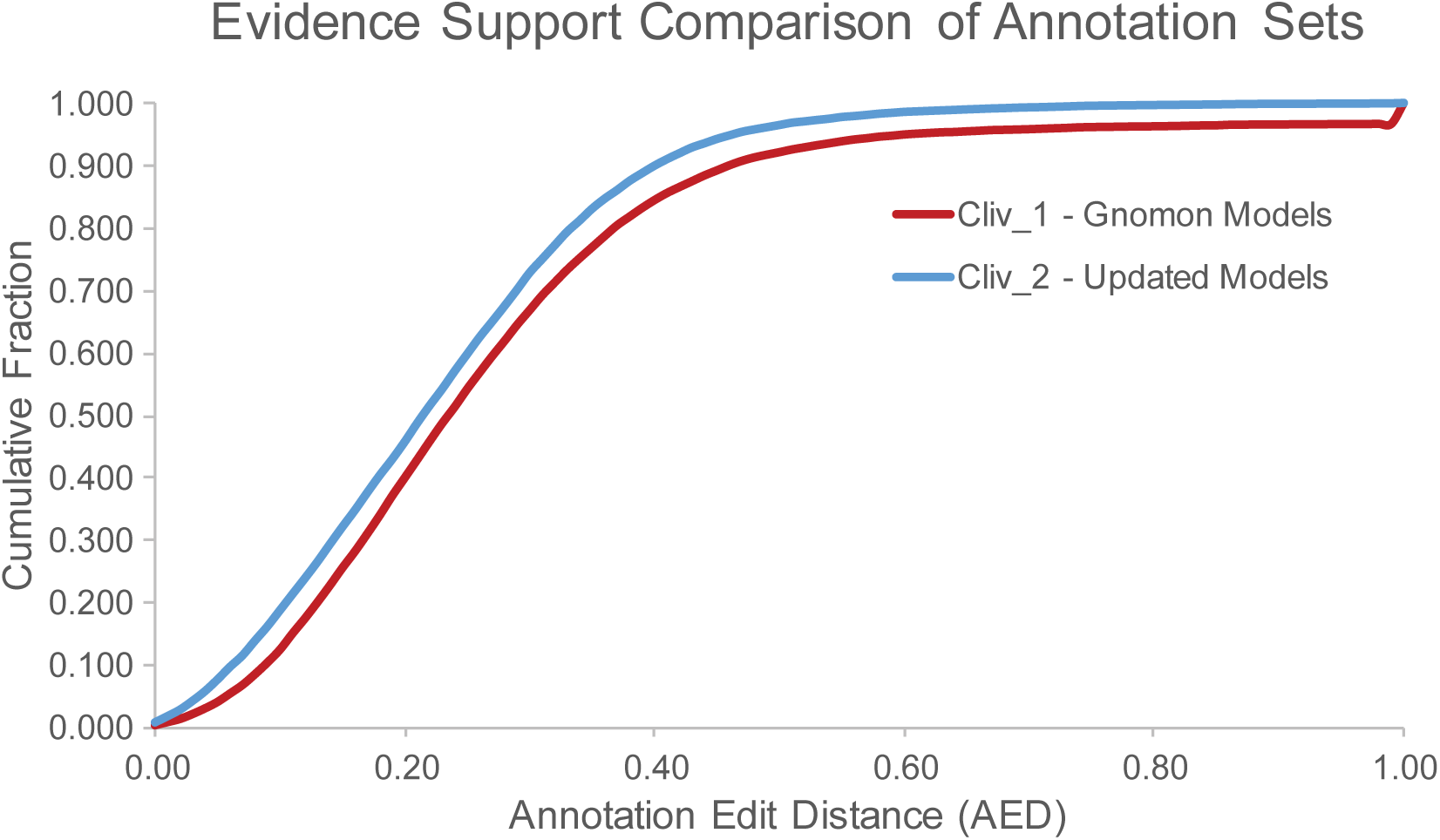
Evidence support comparison of annotation sets. Annotation edit distance (AED) support for gene models in Cliv_2.1 (blue line) is improved over Cliv_1.0 (NCBI Gnomon annotation, red line).

**Figure 4.**
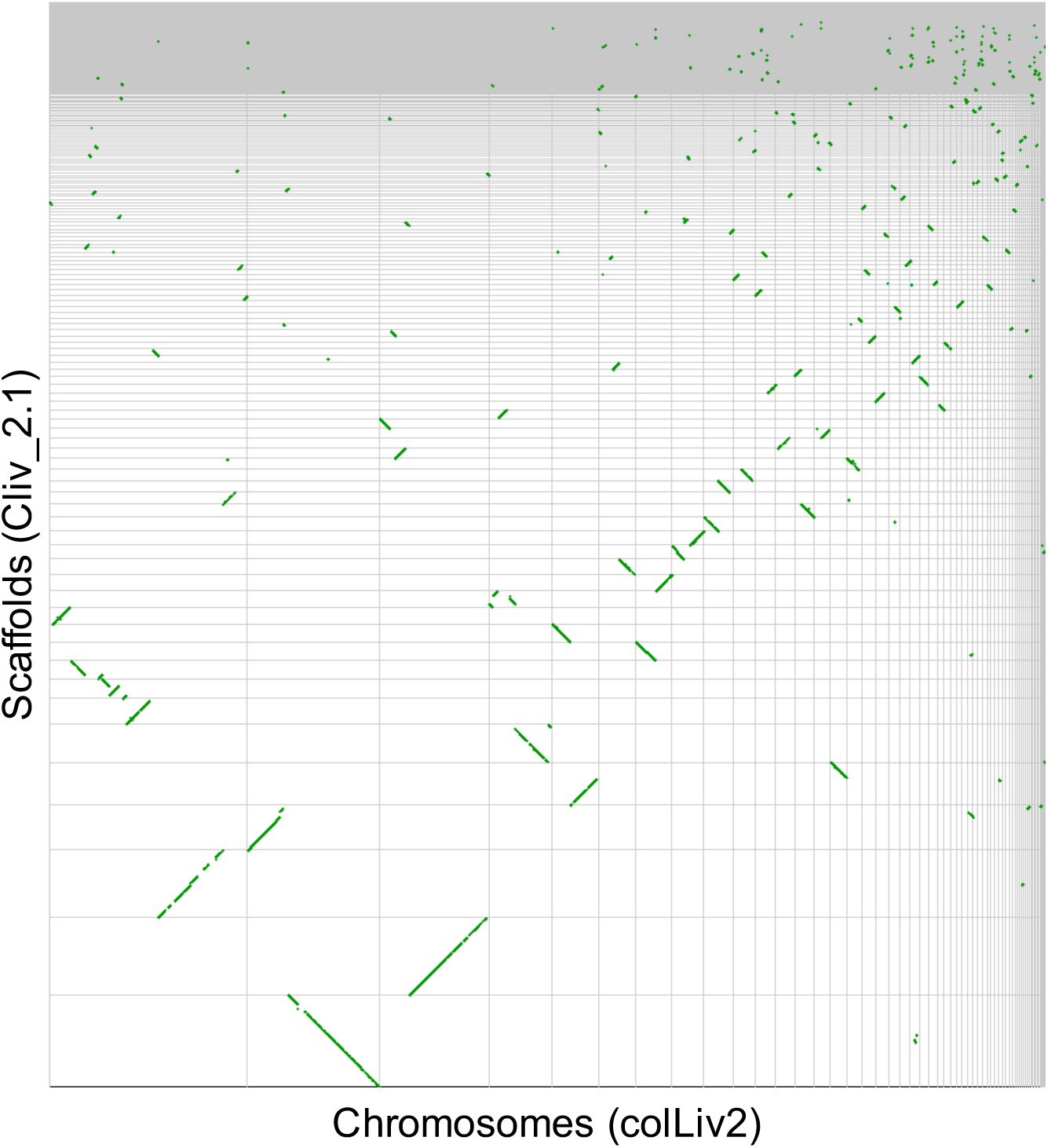
Dot plot of syntenic regions between the Cliv_2.1 and colLiv2 assemblies of the *C. livia* genome. Each segment of the X axis represents a single colLiv2 scaffold ordered from largest (left) to smallest (right), while each segment of the Y axis represents a scaffold of the Cliv_2.1 assembly, ordered from largest (bottom) to smallest (top). Green dots indicate aligned regions of synteny.

### Linkage map

The linkage map consists of 3,192 markers assembled into 48 autosomal linkage groups and a single Z-chromosome linkage group (Table S6. The map contains markers from 236 scaffolds. Together, these scaffolds encompass 1,048,536,443 bp (94.6%) of the Cliv_2.1 assembly, and include 13,026 of 15,392 annotated genes. Cliv_2.1 scaffolds are strongly supported by linkage data. For 235 out of 236 scaffolds included in the linkage map, all GBS markers mapped to that scaffold form a single contiguous block within one linkage group (scaffold ScoHet5_252 was split between two linkage groups). Additionally, within-scaffold marker order was largely supported by calculated pairwise recombination fractions.

### Comparison with colLiv2 genome assembly

Recently, Damas et al. (2017) used computational methods and universal avian bacterial artificial chromosome (BAC) probes to achieve chromosome-level scaffolding using the Cliv_1.0 assembly as input material. This assembly, named colLiv2 (GenBank assembly accession GCA_001887795.1; 1,018,016,946 bp in length), is approximately 8% smaller than the Cliv_2.1 assembly.

Based on genome-wide pairwise alignments using LAST (Fig. 4)(Kielbasa et al. 2011), a substantial number of regions of Cliv_2.1 that do not align to colLiv2 genome contain both unique sequence and annotated genes. Based on gene coordinates, 1184 annotated Cliv_2.1 genes were absent from colLiv2 (Table 6).

**Table 6.**
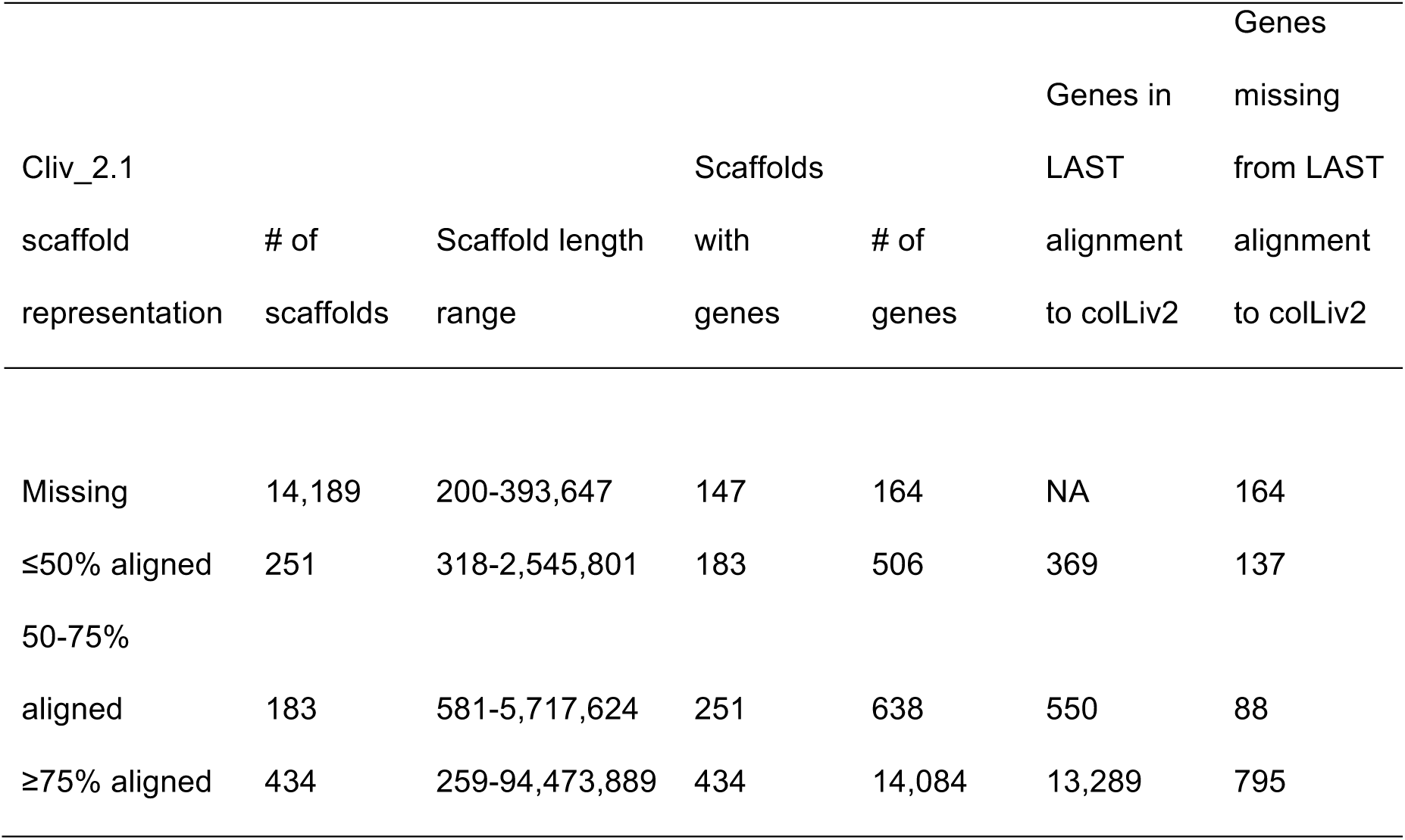
Summary of Cliv_2.1 alignment to colLiv2 chromosome-level scaffolds. Overall, colLiv2 appears to exclude 1,184, or approximately 7.7%, of the 15,392 annotated genes from the Cliv_2.1 assembly; this is consistent with the overall decrease in genome size.

Of the 3,192 GBS makers mapped to Cliv_2.1, 2,940 markers (92.1%) mapped to colLiv2 with an E-value <4e-24. Of the remaining markers, 7 mapped to colLiv2 with an E-value >4e-24, and 245 markers (7.67%) failed to map to colLiv2 entirely. We assessed the agreement between marker and linkage data by calculating pairwise recombination fractions for the 2940 markers, then plotted these recombination fractions in the order in which markers appear on the colLiv2 chromosome-level scaffolds. Overall, the marker order largely agrees with calculated recombination fractions; however, we identified a number of locations where pairwise recombination fractions suggest that portions of the colLiv2 chromosomes are not ordered properly, as exemplified in Fig. 5. We also identified 42 markers for which the location with the best sequence match in colLiv2 appears to be incorrect based on recombination fraction estimates; these markers are summarized in Table S7.

**Figure 5.**
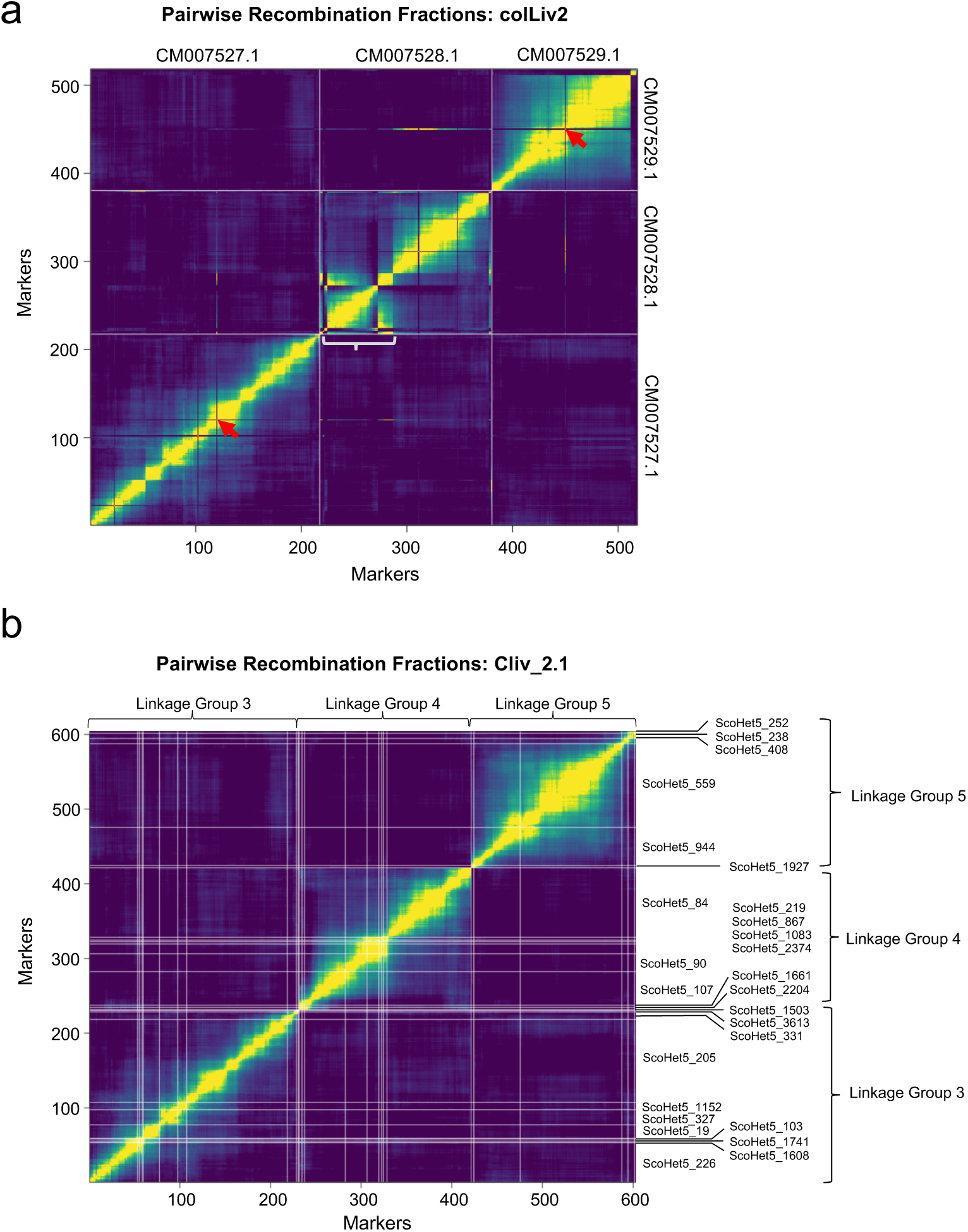
Correspondence between genotyping data and marker order in colLiv2 and Cliv_2.1 assemblies. (a) Representative plot of pairwise recombination fractions for GBS markers, ordered based on best alignment to colLiv2 assembly, for chromosomes CM007527.1, CM007528.1, and CM007529.1. X and Y axes show individual markers, ordered as they map to the colLiv2 chromosomes CM007527.1, CM007528.1, and CM007529.1. White lines mark the boundaries between chromosomes. Yellow indicates low pairwise recombination fraction (linked markers), while purple indicates high pairwise recombination fraction (unlinked markers). Red arrows highlight two markers, one mapped to chromosome CM007527.1 and one mapped to CM007529.1, for which recombination fractions suggest that these markers should instead be located on chromosome CM007528.1. A white bracket indicates a region on chromosome CM007528.1 where portions of the chromosome appear to be assembled in the wrong order. (b) Plot of pairwise recombination fractions for the Cliv_2.1 scaffolds that make up linkage groups 3, 4, and 5. In (a), colLiv2 CM007527.1 largely corresponds to linkage group 3, CM007528.1 to linkage group 4, and CM007529.1 to linkage group 5. White lines mark the boundaries between individual scaffolds, with scaffold IDs indicated on the right side.

### Conclusions

The improved scaffold lengths and updated gene model annotations of Cliv_2.1 will further empower ongoing studies to identify genes responsible for phenotypic traits of interest. In addition, longer scaffolds will improve detection of regions under selection, including large deletions and other structural variants responsible for interesting traits in *C. livia*. Finally, our new transcriptomic data provide tissue-specific expression profiles for several adult tissue types and an important embryonic stage for the morphogenesis of limbs, craniofacial structures, skin, and other tissues.

## ACKNOWLEDGEMENTS

We thank Dovetail Genomics for their aid in scaffolding the assembly; Julia Carleton and Anna Vickrey for technical support; and Elena Boer for comments on the manuscript. This work was supported by National Science Foundation grant DEB1149160 and National Institutes of Health (NIH) grant R01GM115996 to MDS; NSF EAGER grant IOS1561337 to MY; a European Research Council starting grant 336724 and Austrian Science Fund (FWF) grant Y726 to DAK; and European Research Council Consolidator grant 681396 to MTPG. We gratefully acknowledge research support from Boehringer Ingelheim at the Research Institute of Molecular Pathology, and support and resources from the Center for High Performance Computing at the University of Utah.

## SUPPLEMENTAL TABLES

**Table S1. Positions of Cliv_1.0 scaffolds in the Cliv_2.1 scaffolds.** The table has the following format: column 1, Cliv_2.1 scaffold name; column 2, Cliv_1.0 sequence name; column 3, starting base (zero-based) of the Cliv_1.0 sequence; column 4, ending base of the Cliv_1.0 sequence; column 5, orientation of the Cliv_1.0 sequence in the Cliv_2.1 scaffold, where (-) indicates that the Cliv_2.1 scaffold sequence is reverse complemented relative to the Cliv_1.0 assembly; column 6, starting base (zero-based) in the Cliv_2.1 scaffold; column 7, ending base in the Cliv_2.1 scaffold.

**Table S2. Positions of breaks made in the Cliv_1.0 assembly to create the Cliv_2.1 assembly**. Data fields follow the same format that is used in Supplemental Table 1.

**Table S3. Summary of transposable element fragments, parsed into 1 My bins based on substitution rate**

**Table S4. Summary information of repeat masking, by class and by family**

**Table S5. Transcript count and cumulative distribution function (CDF) binned by Annotation Edit Distance (AED) values**. AED is a modified sensitivity/specificity metric used to compare annotation datasets to each other or to aligned transcriptome and protein homology datasets. For calculating AED, sensitivity is defined as the fraction of a given reference overlapping a prediction and measures false negative rates. For our purposes, the prediction is a transcript model and the reference (or truth set) is a set of aligned transcriptome and protein homology evidence. We calculate sensitivity using the formula SN = |p∩r|/|r|; where |p∩r| represents the number overlapping nucleotides between the prediction and reference, and |r| represents the total number of nucleotides in the reference. Specificity is then defined as the fraction of a prediction overlapping a given reference, and it measures false positive rates. We calculate specificity using the formula SP = |p∩r|/|p|. We then define concordance to be the average of sensitivity and specificity (C = (SN+SP)/2), and AED is 1 minus the concordance (AED = 1- C). Transcript models that have high AED values then show little concordance to aligned experimental evidence, and models with low AED values show high concordance.

**Table S6. Linkage map assembled from genotype-by-sequencing markers aligned to the Cliv_2.1 assembly, and positions of aligned markers within the Cliv_2.1, Cliv_1.0, and colLiv2 assemblies**.

**Table S7. Summary of GBS markers for which best BLAST alignment to colLiv2 is discordant with linkage data.** Columns describe marker position in the linkage map, the best BLAST hit within the colLiv2 assembly, and the marker position in the Cliv_2.1 assembly. For each marker, the colLiv2 chromosome to which the marker appears to be linked is also indicated.

## REFERENCES

Altschul, S. F., W. Gish, W. Miller, E. W. Meyers, and D. J. Lipman. 1990. Basic Local Alignment Search Tool. Journal of Molecular Biology 215:403–410.

Bairoch, A. and R. Apweiler. 2000. The SWISS-PROT protein sequence database and its supplement TrEMBL in 2000. Nucl. Acids Res. 28:45–48.

Bao, W., K. K. Kojima, and O. Kohany. 2015. Repbase Update, a database of repetitive elements in eukaryotic genomes. Mob DNA 6:11.

Broman, K., H. Wu, S. Sen, and G. Churchill. 2003. R/qtl: QTL mapping in experimental crosses. Bioinformatics 19:889–890.

Cantarel, B. L., I. Korf, S. M. C. Robb, G. Parra, E. Ross, B. Moore, C. Holt, A. Sanchez Alvarado, and M. Yandell. 2008. MAKER: An easy-to-use annotation pipeline designed for emerging model organism genomes. Genome Res. 18:188–196.

Catchen, J. M., A. Amores, P. Hohenlohe, W. Cresko, and J. H. Postlethwait. 2011. Stacks: building and genotyping loci de novo from short-read sequences. G3 1:171–182.

Damas, J., R. O’Connor, M. Farre, V. P. E. Lenis, H. J. Martell, A. Mandawala, K. Fowler,S. Joseph, M. T. Swain, D. K. Griffin, and D. M. Larkin. 2017. Upgrading short-read animal genome assemblies to chromosome level using comparative genomics and a universal probe set. Genome Res 27:875–884.

Domyan, E. T., M. W. Guernsey, Z. Kronenberg, S. Krishnan, R. E. Boissy, A. I. Vickrey,C. Rodgers, P. Cassidy, S. A. Leachman, J. W. Fondon, 3rd, M. Yandell, and M. D. Shapiro. 2014. Epistatic and combinatorial effects of pigmentary gene mutations in the domestic pigeon. Curr Biol 24:459–464.

Domyan, E. T., Z. Kronenberg, C. R. Infante, A. I. Vickrey, S. A. Stringham, R. Bruders,M. W. Guernsey, S. Park, J. Payne, R. B. Beckstead, G. Kardon, D. B. Menke, M. Yandell, and M. D. Shapiro. 2016. Molecular shifts in limb identity underlie development of feathered feet in two domestic avian species. eLife 5:e12115.

Domyan, E. T. and M. D. Shapiro. 2017. Pigeonetics takes flight: Evolution, development, and genetics of intraspecific variation. Dev Biol 427:241–250.

Eilbeck, K., B. Moore, C. Holt, and M. Yandell. 2009. Quantitative measures for the management and comparison of annotated genomes. BMC Bioinformatics 10:67.

Feschotte, C., U. Keswani, N. Ranganathan, M. L. Guibotsy, and D. Levine. 2009. Exploring repetitive DNA landscapes using REPCLASS, a tool that automates the classification of transposable elements in eukaryotic genomes. Genome Biol Evol 1:205–220.

Grabherr, M. G., B. J. Haas, M. Yassour, J. Z. Levin, D. A. Thompson, I. Amit, X. Adiconis,L. Fan, R. Raychowdhury, Q. Zeng, Z. Chen, E. Mauceli, N. Hacohen, A. Gnirke,N. Rhind, F. di Palma, B. W. Birren, C. Nusbaum, K. Lindblad-Toh, N. Friedman, and A. Regev. 2011. Full-length transcriptome assembly from RNA-Seq data without a reference genome. Nature biotechnology 29:644–652.

Hamburger, V. and H. L. Hamilton. 1951. A series of normal stages in the development of the chick embryo. Journal of Morphology 88:49–92.

Holt, C. and M. Yandell. 2011. MAKER2: an annotation pipeline and genome- database management tool for second-generation genome projects. BMC Bioinformatics 12:491.

International Chicken Genome Sequencing, C. 2004. Sequence and comparative analysis of the chicken genome provide unique perspectives on vertebrate evolution. Nature 432:695–716.

Kapusta, A. and A. Suh. 2017. Evolution of bird genomes-a transposon’s-eye view. Ann N Y Acad Sci 1389:164–185.

Kapusta, A., A. Suh, and C. Feschotte. 2017. Dynamics of genome size evolution in birds and mammals. Proc Natl Acad Sci U S A 114:E1460–E1469.

Kielbasa, S. M., R. Wan, K. Sato, P. Horton, and M. C. Frith. 2011. Adaptive seeds tame genomic sequence comparison. Genome Res 21:487–493.

Kronenberg, Z. N., E. J. Osborne, K. R. Cone, B. J. Kennedy, E. T. Domyan, M. D. Shapiro, N. C. Elde, and M. Yandell. 2015. Wham: Identifying Structural Variants of Biological Consequence. PLoS Comput Biol 11:e1004572.

Langmead, B. and S. L. Salzberg. 2012. Fast gapped-read alignment with Bowtie 2. Nat Methods 9:357–359.

Levi, W. M. 1986. The Pigeon (Second Revised Edition). Levi Publishing Co., Inc.,

Sumter, S.C. Lincoln, S. E. and E. S. Lander. 1992. Systematic detection of errors in genetic linkage data. Genomics 14:604–610.

Lyons, E. and M. Freeling. 2008. How to usefully compare homologous plant genes and chromosomes as DNA sequences. Plant J 53:661–673.

Lyons, E., B. Pedersen, J. Kane, M. Alam, R. Ming, H. Tang, X. Wang, J. Bowers, A. Paterson, D. Lisch, and M. Freeling. 2008. Finding and comparing syntenic regions among Arabidopsis and the outgroups papaya, poplar, and grape: CoGe with rosids. Plant physiology 148:1772–1781.

Miller, S. A., D. D. Dykes, and H. F. Polesky. 1988. A simple salting out procedure for extracting DNA from human nucleated cells. Nucleic Acids Res 16:1215.

Morgulis, A., E. M. Gertz, A. A. Schaffer, and R. Agarwala. 2006. WindowMasker: window-based masker for sequenced genomes. Bioinformatics 22:134–141.

Price, A. L., N. C. Jones, and P. A. Pevzner. 2005. De novo identification of repeat families in large genomes. Bioinformatics 21 Suppl 1:i351–358.

Pruitt, K. D., T. Tatusova, and D. R. Maglott. 2007. NCBI reference sequences (RefSeq): a curated non-redundant sequence database of genomes, transcripts and proteins. Nucleic Acids Res:D61 – 65.

Putnam, N. H., B. L. O’Connell, J. C. Stites, B. J. Rice, M. Blanchette, R. Calef, C. J. Troll,A. Fields, P. D. Hartley, C. W. Sugnet, D. Haussler, D. S. Rokhsar, and R. E. Green. 2016. Chromosome-scale shotgun assembly using an in vitro method for long-range linkage. Genome Res 26:342–350.

Shapiro, M. D., Z. Kronenberg, C. Li, E. T. Domyan, H. Pan, M. Campbell, H. Tan, C. D. Huff, H. Hu, A. I. Vickrey, S. C. Nielsen, S. A. Stringham, H. Hu, E. Willerslev, M.T. Gilbert, M. Yandell, G. Zhang, and J. Wang. 2013. Genomic diversity and evolution of the head crest in the rock pigeon. Science 339:1063–1067.

Simao, F. A., R. M. Waterhouse, P. Ioannidis, E. V. Kriventseva, and E. M. Zdobnov. 2015. BUSCO: assessing genome assembly and annotation completeness with single-copy orthologs. Bioinformatics 31:3210–3212.

Slater, G. and E. Birney. 2005. Automated generation of heuristics for biological sequence comparison. BMC Bioinformatics 6:31.

Smit, A. F. and R. Hubley. 2008. RepeatModeler Open-1.0 http://www.repeatmasker.org/.

Smit, A. F., R. Hubley, and P. Green. 1996. RepeatMasker Open-3.0 http://www.repeatmasker.org/.

Smit, A. F., R. Hubley, and P. Green. 2015. RepeatMasker Open-4.0.2013-2015 http://www.repeatmasker.org/.

Souvorov, A., Y. Kapustin, B. Kiryutin, V. Chetvernin, T. Tatusova, and D. Lipman. 2010. Gnomon – NCBI eukaryotic gene prediction tool. NCBI.

Stanke, M., M. Diekhans, R. Baertsch, and D. Haussler. 2008. Using native and syntenically mapped cDNA alignments to improve de novo gene finding. Bioinformatics 24:637–644.

Stanke, M. and S. Waack. 2003. Gene prediction with a hidden Markov model and a new intron submodel. Bioinformatics 19:ii215–225.

Suh, A., C. C. Witt, J. Menger, K. R. Sadanandan, L. Podsiadlowski, M. Gerth, A. Weigert, J. A. McGuire, J. Mudge, S. V. Edwards, and F. E. Rheindt. 2016. Ancient horizontal transfers of retrotransposons between birds and ancestors of human pathogenic nematodes. Nature communications 7:11396.

UniProt, C. 2007. The Universal Protein Resource (UniProt). Nucleic Acids Res:D193 – 197.

Warren, W. C., D. F. Clayton, H. Ellegren, A. P. Arnold, L. W. Hillier, A. Kunstner, S. Searle, S. White, A. J. Vilella, S. Fairley, A. Heger, L. Kong, C. P. Ponting, E. D. Jarvis, C. V. Mello, P. Minx, P. Lovell, T. A. Velho, M. Ferris, C. N. Balakrishnan,S. Sinha, C. Blatti, S. E. London, Y. Li, Y. C. Lin, J. George, J. Sweedler, B. Southey, P. Gunaratne, M. Watson, K. Nam, N. Backstrom, L. Smeds, B. Nabholz, Y. Itoh, O. Whitney, A. R. Pfenning, J. Howard, M. Volker, B. M. Skinner, D. K. Griffin, L. Ye, W. M. McLaren, P. Flicek, V. Quesada, G. Velasco, C. Lopez-Otin, X. S. Puente, T. Olender, D. Lancet, A. F. Smit, R. Hubley, M. K. Konkel, J. A. Walker, M. A. Batzer, W. Gu, D. D. Pollock, L. Chen, Z. Cheng, E. E. Eichler, J. Stapley, J. Slate, R. Ekblom, T. Birkhead, T. Burke, D. Burt, C. Scharff,I. Adam, H. Richard, M. Sultan, A. Soldatov, H. Lehrach, S. V. Edwards, S. P. Yang, X. Li, T. Graves, L. Fulton, J. Nelson, A. Chinwalla, S. Hou, E. R. Mardis, and R. K. Wilson. 2010. The genome of a songbird. Nature 464:757–762.

Yokouchi, Y., M. Yamamoto, T. Toyota, H. Sasaki, and A. Kuroiwa. 1993. Regulatory interaction of positional signalings on coordinate expression of homeobox genes in developing limb buds. Limb Development and Regeneration. Wiley- Liss, Inc.

Zhang, G., B. Li, C. Li, M. T. Gilbert, E. D. Jarvis, J. Wang, and C. Avian Genome. 2014a. Comparative genomic data of the Avian Phylogenomics Project. Gigascience 3:26.

Zhang, G., C. Li, Q. Li, B. Li, D. M. Larkin, C. Lee, J. F. Storz, A. Antunes, M. J. Greenwold,R. W. Meredith, A. Odeen, J. Cui, Q. Zhou, L. Xu, H. Pan, Z. Wang, L. Jin, P. Zhang, H. Hu, W. Yang, J. Hu, J. Xiao, Z. Yang, Y. Liu, Q. Xie, H. Yu, J. Lian, P. Wen, F. Zhang, H. Li, Y. Zeng, Z. Xiong, S. Liu, L. Zhou, Z. Huang, N. An, J. Wang,Q. Zheng, Y. Xiong, G. Wang, B. Wang, J. Wang, Y. Fan, R. R. da Fonseca, A. Alfaro-Nunez, M. Schubert, L. Orlando, T. Mourier, J. T. Howard, G. Ganapathy, A. Pfenning, O. Whitney, M. V. Rivas, E. Hara, J. Smith, M. Farre, J. Narayan, G. Slavov, M. N. Romanov, R. Borges, J. P. Machado, I. Khan, M. S. Springer, J. Gatesy, F. G. Hoffmann, J. C. Opazo, O. Hastad, R. H. Sawyer, H. Kim, K. W. Kim,H. J. Kim, S. Cho, N. Li, Y. Huang, M. W. Bruford, X. Zhan, A. Dixon, M. F. Bertelsen, E. Derryberry, W. Warren, R. K. Wilson, S. Li, D. A. Ray, R. E. Green, S. J. O’Brien, D. Griffin, W. E. Johnson, D. Haussler, O. A. Ryder, E. Willerslev, G.R. Graves, P. Alstrom, J. Fjeldsa, D. P. Mindell, S. V. Edwards, E. L. Braun, C. Rahbek, D. W. Burt, P. Houde, Y. Zhang, H. Yang, J. Wang, E. D. Jarvis, M. T. Gilbert and J. Wang. 2014b. Comparative genomics reveals insights into avian genome evolution and adaptation. Science 346:1311–1320.

